# *De novo EIF2AK1* and *EIF2AK2* variants are associated with developmental delay, leukoencephalopathy, and neurologic decompensation

**DOI:** 10.1101/757039

**Authors:** Dongxue Mao, Chloe M. Reuter, Maura R.Z. Ruzhnikov, Anita E. Beck, Emily G. Farrow, Lisa T. Emrick, Jill A. Rosenfeld, Katherine M. Mackenzie, Laurie Robak, Matthew T. Wheeler, Lindsay C. Burrage, Mahim Jain, Pengfei Liu, Daniel Calame, Sebastien Küry, Martin Sillesen, Klaus Schmitz-Abe, Davide Tonduti, Luigina Spaccini, Maria Iascone, Casie A. Genetti, Madeline Graf, Alyssa Tran, Mercedes Alejandro, Undiagnosed Diseases Network, Brendan H. Lee, Isabelle Thiffault, Pankaj B. Agrawal, Jonathan A. Bernstein, Hugo J. Bellen, Hsiao-Tuan Chao

## Abstract

*EIF2AK1* and *EIF2AK2* encode members of the Eukaryotic Translation Initiation Factor 2 Alpha Kinase (EIF2AK) family that inhibits protein synthesis in response to physiologic stress conditions. EIF2AK2 is also involved in innate immune response and the regulation of signal transduction, apoptosis, cell proliferation, and differentiation. Despite these findings, human disorders associated with deleterious variants in *EIF2AK1* and *EIF2AK2* have not been reported. Here, we describe the identification of eight unrelated individuals with heterozygous *de novo* missense variants in *EIF2AK1* (1/8) or *EIF2AK2* (7/8). Features seen in these eight individuals include white matter alterations (8/8), developmental delay (8/8), impaired language (8/8), cognitive impairment (7/8), ataxia (6/8), dysarthria in probands with verbal ability (6/6), hypotonia (6/8), hypertonia (5/8), and involuntary movements (3/8). Individuals with *EIF2AK2* variants also exhibit neurological regression in the setting of febrile illness or infection. We use mammalian cell lines and patient-derived fibroblasts to further confirm the pathogenicity of variants in these genes and found reduced kinase activity. EIF2AKs phosphorylate Eukaryotic Translation Initiation Factor 2 Subunit 1, (EIF2S1, also known as EIF2α), which then inhibits EIF2B activity. Deleterious variants in genes encoding EIF2B proteins cause childhood ataxia with central nervous system hypomyelination/vanishing white matter disease (CACH/VWM), a leukoencephalopathy characterized by neurologic regression in the setting of febrile illness and other stressors. Our findings indicate that *EIF2AK2* missense variants cause a neurodevelopmental syndrome that may share phenotypic and pathogenic mechanisms with CACH/VWM.

## REPORT

The Eukaryotic Translation Initiation Factor 2 Alpha Kinase (EIF2AK) family is comprised of four mammalian kinases that regulate the cytoprotective integrated stress response (ISR) required for cellular adaptation to stress conditions^1; 2^. EIF2AK1 (MIM: *613635; HGNC: 24921), also known as Heme-Regulated Inhibitor, responds to heme deprivation, proteasome inhibition, and maintains basal endoplasmic reticulum (ER) stress^3–7^. EIF2AK1 contains two protein kinase domains and two heme binding sites. EIF2AK2 (MIM: *176871; HGNC: 9437), also known as Protein Kinase R (PKR), is activated by double-stranded RNA (dsRNA) and can block the translation of viral mRNA in response to infection^8–10^, activation also occurs in response to oxidative stress, ER stress^11–14^, cytokines^14; 15^, and growth factors^16^. EIF2AK2 contains two dsRNA binding motifs (DSRM) and a protein kinase domain. In the presence of their respective cellular stressors, both EIF2AK1 or EIF2AK2 activate ISR by phosphorylating Eukaryotic Translation Initiation Factor 2 Subunit 1 (EIF2S1, also known as EIF2α), a major regulator of the initiation of mRNA translation and the rate of protein synthesis. Phosphorylation of EIF2S1 on serine 51 by EIF2AK family members prevents mRNA translation and result in transient suppression of general protein synthesis^17; 18^. Prior studies have linked missense, nonsense, and splicing variants in *EIF2AK3* and truncating variants in *EIF2AK4* to autosomal recessive epiphyseal dysplasia with early onset diabetes mellitus (OMIM #226980) and autosomal recessive pulmonary veno-occlusive disease type 2 (OMIM #234810), respectively^19; 20^. Neither of these disorders present with primary neurologic findings. However, the phenotypic consequences of rare variants in human *EIF2AK1* and *EIF2AK2* are currently unknown.

Through the Undiagnosed Diseases Network^25; 26^, curation of ~13,500 clinical exome sequencing (ES) from Baylor Genetics (BG), and GeneMatcher^21; 22^, we identified multiple unrelated individuals with consistent clinical symptoms, abnormal magnetic resonance imaging (MRI) findings in the brain and/or spinal cord (**Figure 1A-F**), and rare *de novo* missense variants in either *EIF2AK1* or *EIF2AK2*. Eight individuals were enrolled into our study. All had rare missense variants in either *EIF2AK1* or *EIF2AK2* and were identified via trio ES with Sanger sequencing confirmation. The variants localize to the protein kinase or DSRM domains of EIF2AK1 and EIF2AK2^9–11^ (**Fig2A, B**). Additionally, we identified a rare missense variant in *EIF2AK2* (c.341T>A, p.Leu114Gln) of unknown inheritance from a proband-only ES in a patient with discordant phenotypes. We included the *EIF2AK2* p.Leu114Gln variant in the molecular studies as a rare variant control. All of the variants were absent from the Exome Aggregation Consortium Database^23^ and the Genome Aggregation Database (gnomAD)^24; 27^.

**Figure 1:**
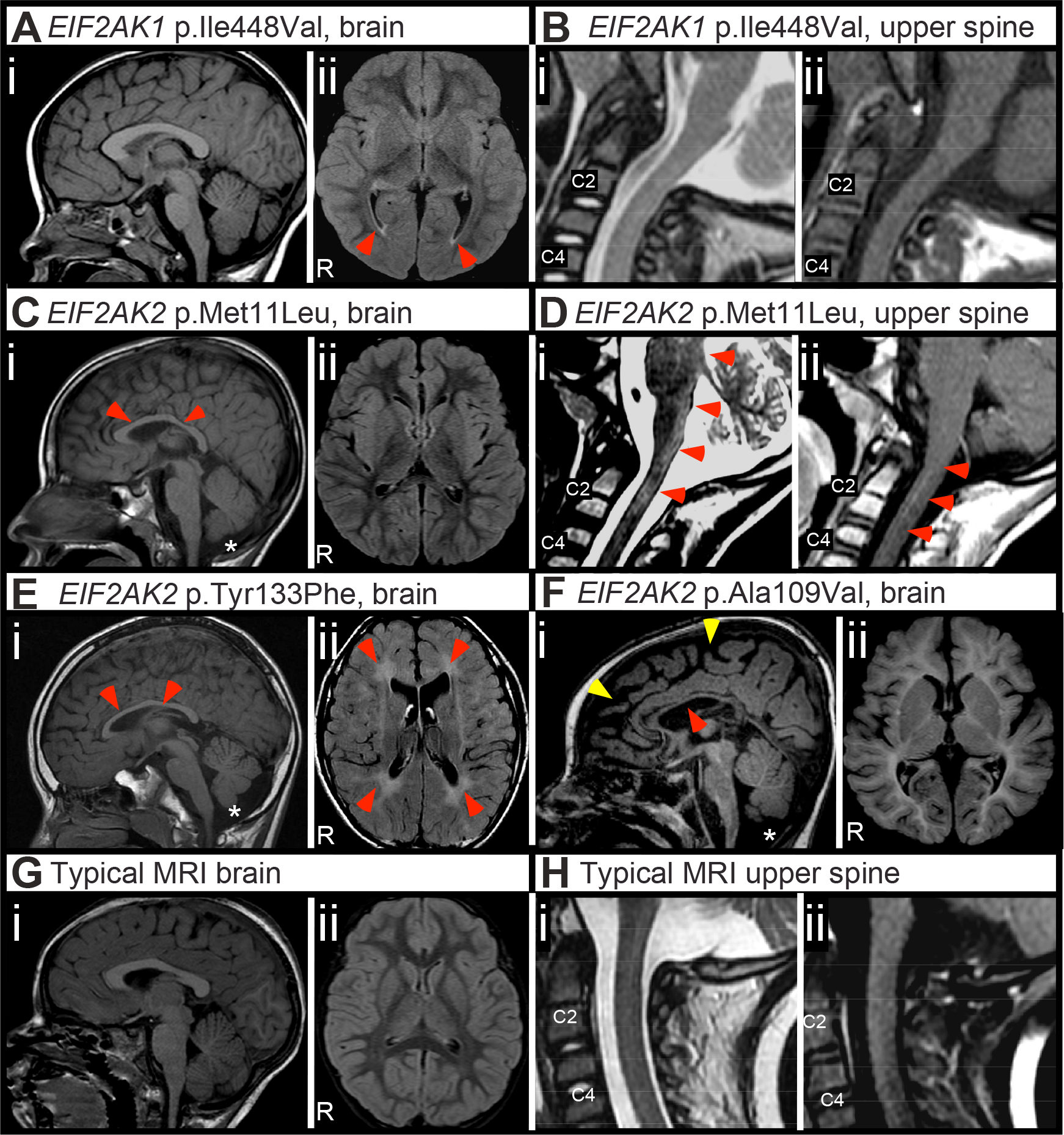
Hypomyelination, cerebral atrophy, and white matter abnormalities associated with *de novo EIF2AK2* missense variants. **(A, B)** Representative images from proband 1 with *EIF2AK1* p.Ile448Val variant acquired on a 1.5 Tesla (1.5T) MRI. **(A)** MRI brain without contrast images. **(i)** Mid-sagittal T1-weighted image with appropriate size of corpus callosum and cerebellar vermis. **(ii)** Axial FLAIR image showing T2-weighted hyperintensities at the posterior lateral ventricles (red arrowheads). **(B)** MRI upper spinal cord images. **(i)** Mid-sagittal T2-weighted and **(ii)** T1-weighted images showing unremarkable upper spinal cord appearance. **(C, D)** Representative images from proband 2 with *EIF2AK2* p.Met11Leu variant acquired on a 1.5 Tesla (1.5T) MRI. **(C)** MRI brain without contrast images. **(i)** Mid-sagittal T1-weighted image showing thinning of the corpus callosum (red arrowheads) and mild cerebellar vermis hypoplasia (asterisk). **(ii)** Axial FLAIR image showing mild reduction in cerebral volume with thinning of the gyri and widening of the sulci. **(D)** MRI upper spinal cord images. **(i)** Mid-sagittal T2-weighted image showing hyperintensities in the dorsal-most upper cervical cord, dorsal medulla, and dorsal pons (red arrowheads) **(ii)** Post-contrast mid-sagittal T1-weighted image showing contrast enhancement in the upper cervical cord. **(E)** Representative images from proband 3 with *EIF2AK2* p.Tyr133Phe variant acquired on a 1.5T MRI. **(i)** Mid-sagittal T1-weighted image showing thinning of the corpus callosum (red arrowheads) with cerebellar vermis hypoplasia and prominent cisterna magna (asterisk). **(ii)** Axial FLAIR image showing diffuse hyperintensities throughout the white matter (red arrowheads). **(F)** Representative images from proband 6 with *EIF2AK2* p.Ala109Val acquired on a 3.0T MRI. **(i)** Mid-sagittal T1-weighted image showing thinning of the corpus callosum (red arrowhead), cerebral atrophy (yellow arrowheads), and cerebellar vermis hypoplasia (asterisk). Axial FLAIR image showing pronounced hypomyelination in the cerebral hemispheres. **(G)** Representative typical control MRI brain **(i)** mid-sagittal T1-weighted and **(ii)** axial FLAIR images acquired on a 1.5T MRI. **(H)** Representative typical control **(i)** mid-sagittal T2-weighted and **(ii)** post-contrast mid-sagittal T1-weighted images acquired on a 1.5T MRI.

**Figure 2:**
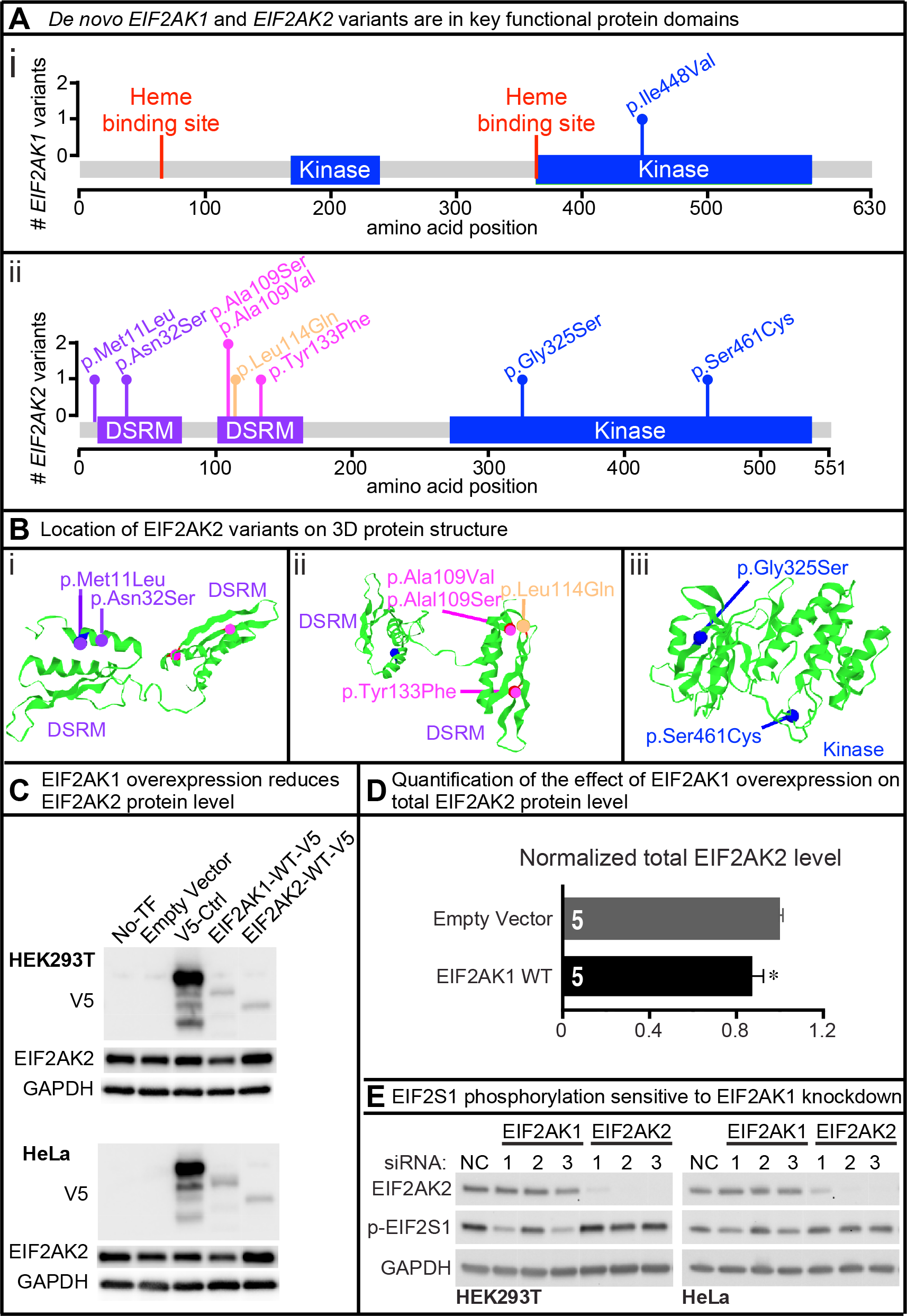
*De novo EIF2AK1* and *EIF2AK2* missense variants map to key protein domains and *EIF2AK1* p.Ile448Val impairs kinase activity. **(A)** Lollipop plots showing variants relative to a schematic representation of the gene adapted from MutationMapper. Heme-binding sites in red, protein kinase domain (Kinase) in blue and double-stranded RNA-binding motif (DSRM) in purple. **(i)** *EIF2AK1* variant is located in the kinase domain and **(ii)** *EIF2AK2* variants are located in the DSRM and Kinase domains. **(B)** 3D structure of EIF2AK2 DSRM and Kinase domains with d*e novo EIF2AK2* variants in purple, magenta, or blue. The rare variant control, p.Leu114Gln, is in orange. Variants are mapped to the protein 3D structure using Mutation3D^48^. Protein Data Bank ID: 1QU6 and 3UIU. **(C)** Full length human *EIF2AK1, EIF2AK2*, and unrelated control cDNAs were cloned into pcDNA™-DEST40 Vector with a CMV promoter and C-terminus V5 tag. Lipofectamine™ 3000 was used to transfect the cDNA vectors into HEK293T and HeLa cells. Western blots show expression level of V5-tagged protein and endogenous level of EIF2AK2. EIF2AK1 overexpression reduces EIF2AK2 protein level in HEK293T and HeLa cell lines. All Western blot images in this paper were acquired using the Bio-Rad ChemiDoc™ Imaging Systems and densitometric analyses of the bands were performed with ImageJ. All images were collected by the imaging system within the linear range. **(D)** Quantification of the effect of EIF2AK1 overexpression in mammalian cell lines on EIF2AK2 protein levels. Statistical significance determined by Student’s t-test. Data shown as mean ± SEM. *n* = 5 independent replicates. * *p* < 0.05 **(E)** Lipofectamine RNAiMAX was used to transfect HEK293T and HeLa cells with either control, *EIF2AK1*, or *EIF2AK2* siRNA for 3 days. Three different siRNAs were tested per gene. Western blots show knockdown efficiency for *EIF2AK1* and *EIF2AK2*. Two EIF2AK1 siRNAs show reduced p-EIF2S1 levels in both HEK293T and HeLa cells. Knockdown of EIF2AK2 does not affect p-EIF2S1 levels in either HEK293T or HeLa cells.

All individuals with *EIF2AK1* or *EIF2AK2* variants had motor delay (8/8), speech and language delay (8/8), and abnormal white matter findings in the central nervous system (8/8). Additional MRI findings include thinning of the corpus callosum (7/8), cerebral volume loss (6/8), diffuse hypomyelination (4/8), and reduced cerebellar vermis volume (4/8). Intellectual function ranges from age appropriate (1/8) to cognitive impairment (7/8). Other features include hypotonia (6/8), hypertonia (5/8), spasticity (6/8), gait and truncal ataxia (6/8), involuntary movements (3/8), and seizures (3/8). Intriguingly, six out of seven individuals (probands 3-8) with *EIF2AK2* variants had neurologic decompensation and loss of developmental milestones with febrile illnesses.

Clinical data were obtained after written informed consent was obtained in accordance with the ethical standards of the participating institutional review boards (IRB) on human research at each respective institution. DNA was extracted from peripheral blood mononuclear cells for ES. The Undiagnosed Diseases Network identified a *de novo* c.1342A>G (p.Ile448Val) variant in *EIF2AK1* (GenBank: NM_014413.4) for proband 1, a *de novo* c.31A>C (p.Met11Leu) variant for proband 2, and a *de novo* c.398A>T (p.Tyr133Phe) variant for proband 3 in *EIF2AK2* (GeneBank: NM_002759.3) by trio ES. Researchers used Codified Genomics (variation interpretation software) for variant review in probands 1 and 2. The Center for Mendelian Genomics and the Broad Institute performed trio ES for proband 5, analyzed the results with SEQR (https://seqr.broadinstitute.org/) and VExP^28^, and identified a *de novo EIF2AK2* c.1382C>G (p.Ser461Cys) variant. Clinically based trio ES, performed in diagnostic labs certified by the Clinical Laboratory Improvement Amendments, identified *de novo EIF2AK2* sequence changes c.973G>A (p.Gly325Ser), c.326C>T (p.Ala109Val), c.325G>T (p.Ala109Ser), and c.95A>G (p.Asn32Ser) in probands 4, 6, 7, and 8, respectively. Maternity and paternity were confirmed by the inheritance of rare SNPs from the parents and sample swap was excluded. There were no pathogenic copy number variants identified in the eight probands by chromosomal microarray. A 47-Mb region with absence of heterozygosity on chromosome 17 (17p11.2q24.1) was identified in proband 6. Chromosome 17 has not been reported with a clinical uniparental disomy phenotype and therefore additional UPD testing was not clinically indicated. The region of AOH would be consistent with the family history of consanguinity.

Statistical models examining observed versus expected functional coding variation reveal that *EIF2AK1* and *EIF2AK2* undergo selective restraint, a process where selection has reduced functional variation, suggesting that missense variants are more likely to be deleterious^24^. Analysis of the observed to the expected loss-of-function (LoF) variation across the genes for *EIF2AK1* and *EIF2AK2* revealed observed/expected (o/e) scores of 0.47 and 0.30, respectively. These o/e results indicate that there is less LoF variation than predicted^24^. Additionally, the Residual Variation Intolerance Score version 4 (RVISv4) is −0.331 for *EIF2AK1* and −1.2108 for *EIF2AK2*, where RIVS < 0 indicates there is less common functional variation in the population than predicted^29^. Finally, both *EIF2AK1* and *EIF2AK2* have low probability of LoF intolerance scores (pLI = 0 and 0.06, respectively) in gnomAD^24^. Based on the gene size and GC content, 34 (*EIF2AK1*) and 32 (*EIF2AK2*) LoF variants were expected, and in the gnomAD population 16 (*EIF2AK1*) and 9 (*EIF2AK2*) LoF variants were observed^24^. Together, these statistical findings indicate that *EIF2AK1* and *EIF2AK2* likely are haplosufficient, but are less tolerant of missense variants.

To determine the functional consequences of the *de novo* variants identified in *EIF2AK1* and *EIF2AK2*, we cloned full-length human wildtype (WT) *EIF2AK1, EIF2AK2*, and unrelated control cDNAs into the mammalian expression vector pcDNA-DEST40. This vector allows for expression of C-terminal V5 (GKPIPNPLLGLDSD) tagged proteins (EIF2AK1-WT-V5 and EIF2AK2-WT-V5) by a CMV promoter. The pcDNA-DEST40 cDNA constructs were transfected into two human cell lines, HEK293T and HeLa. There was a modest level of EIF2AK1-WT-V5 and EIF2AK2-WT-V5 overexpression compared to an unrelated protein-V5 control (**Fig2C**), suggesting that EIF2AK1 and EIF2AK2 protein levels are tightly regulated in these cell lines. Overexpression of EIF2AK1-WT-V5 also reduced the total EIF2AK2 protein level (**Fig2C, D**). Next, to determine the consequences of EIF2AK1 or EIF2AK2 LoF, we examined the impact of either EIF2AK1 or EIF2AK2 knockdown on EIF2S1 phosphorylation (p-EIF2S1) in HEK293T cells or HeLa cells. We designed three independent siRNAs targeting different regions of *EIF2AK1* or *EIF2AK2* mRNA and assessed p-EIF2S1 levels. Two of the three EIF2AK1 siRNAs significantly reduced p-EIF2S1 levels in both HEK293T and HeLa cell lines (**Fig2E**). However, in all three of the EIF2AK2 siRNAs there were no changes in p-EIF2S1 levels (**Fig2E**), suggesting potential redundancy between the EIF2AK family members in HEK293T and HeLa cells.

To test if the *EIF2AK1* and *EIF2AK2* variants are deleterious, we generated pcDNA-DEST40 cDNA constructs to express the human variants in HEK293T or HeLa cells. The variants were generated via either Agilent QuikChange Lightning or NEB Q5 site-directed mutagenesis and confirmed by Sanger sequencing. We assessed the effects of the *EIF2AK1* and *EIF2AK2* variants on protein kinase activity and protein stability in both mammalian cell lines and available patient-derived skin fibroblasts. First, we examined whether the *EIF2AK1* and *EIF2AK2* variants altered protein kinase activity in HEK293T cells. We found that overexpression of EIF2AK1-WT-V5 in HEK293T cells upregulated p-EIF2S1 levels. However, overexpression of EIF2AK1-Ile448Val-V5 had no effect on p-EIF2S1 levels, indicating that the *EIF2AK1* p.Ile448Val variant impairs protein kinase activity (**Fig3A**). Unlike the *EIF2AK1* findings, overexpression of either WT *EIF2AK2* or variants tested in this study had no effect on p-EIF2S1 levels in HEK293T or HeLa cells (**Fig3B, i**). This finding is consistent with our previous observation that EIF2AK2 knockdown in HEK293T and HeLa cells had no effect on p-EIF2S1 levels (**Fig2E**), suggesting that HEK293T and HeLa cells are insensitive to altered EIF2AK2 protein level.

**Figure 3:**
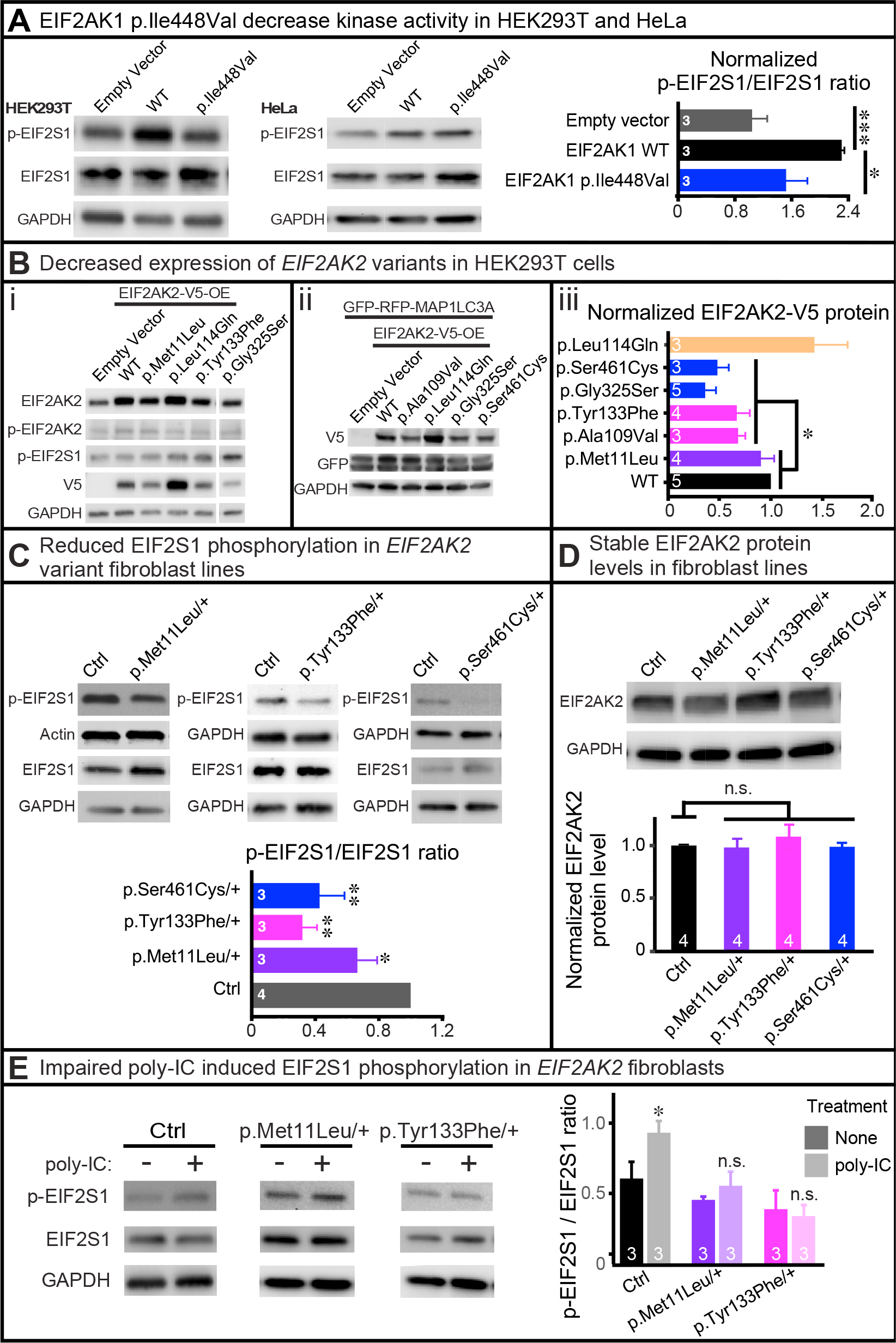
*De novo EIF2AK2* missense variants impair protein stability and kinase activity. **(A)** Lipofectamine™ 3000 was used to transfect HEK293T cells with EIF2AK1-WT-V5 or EIF2AK1-Ile448Val-V5 cDNA vectors. Representative Western blot shows that overexpression of EIF2AK1-Ile448Val-V5 fails to increase EIF2S1 phosphorylation compared to EIF2AK1-WT-V5. Statistical significance determined by confirming the normality of the data (p = 0.05, Shapiro-Wilk test), and then Student’s t-test to measure the difference between groups. Data shown as mean ± SEM. *n* = 3 independent replicates. * *p* < 0.05, ** *p* < 0.01 **(B)** Lipofectamine™ 3000 was used to transfect HEK293T cells with V5-tagged *EIF2AK2* WT or variant cDNAs. Western blots show the expression level of the V5-tagged and endogenous EIF2AK2 protein. **(i)** *EIF2AK2* variants exhibit decreased protein stability compared to WT. No change in EIF2AK2 protein levels were observed with rare variant control, p.Leu114Gln. No change in p-EIF2S1 levels were observed with overexpression of *EIF2AK2* variants. **(ii)** Lipofectamine™ 3000 was used to co-transfect HEK293T cells with GFP-RFP-MAP1LC3A control and either *EIF2AK2*-WT-V5 or *EIF2AK2*-variant-V5 cDNAs. The expression level of GFP is consistent across all cells, indicating that *EIF2AK2* variants do not affect general protein translation. **(iii)** Statistical significance determined by confirming the normality of the data (p = 0.05, Shapiro-Wilk test), and then Student’s t-test to measure the difference between groups. Data shown as mean ± SEM. *n* = 3-5 independent replicates. * *p* < 0.05 **(C)** Western blot showing reduced p-EIF2S1 levels in patient-derived skin fibroblasts with heterozygous *EIF2AK2* missense variants =compared to unrelated control. Statistical significance determined by confirming the normality of the data (p = 0.05, Shapiro-Wilk test), and then Student’s t-test to measure the difference between groups. Data shown as mean ± SEM. *n* = 3-4 independent replicates. * *p* < 0.05, ** *p* < 0.01 **(D)** Western blot showing total EIF2AK2 protein level is not affected in patient-derived skin fibroblasts with heterozygous *EIF2AK2* missense variants compared to unrelated control. Statistical significance determined by confirming the normality of the data (p = 0.05, Shapiro-Wilk test), and then Student’s t-test to measure the difference between groups. *n* = 4 independent replicates. *n.s* = not significant **(E)** Control or patient-derived skin fibroblasts were incubated in regular media with or without poly-IC (final concentration 10 μg/ml) for 24hrs and then protein was collected for Western blot analysis. Increased p-EIF2S1 levels observed in control fibroblasts incubated with poly-IC but patient-derived skin fibroblasts with heterozygous *EIF2AK2* missense variants fail to increase EIF2S1 phosphorylation. Statistical significance determined by Student’s t-test. Data shown as mean ± SEM. *n* = 3 independent replicates. * *p* < 0.05, *n.s* = not significant

Second, we examined whether the *EIF2AK2* variants affected the expression of EIF2AK2 protein in HEK293T (**Fig3B, i-iii**) and HeLa (data not shown) cells by using a V5 antibody to probe for the exogenous protein and an EIF2AK2 antibody to probe for the total EIF2AK2 protein. Interestingly, we found that nearly all *EIF2AK2* variants (p.Tyr133Phe, p.Ala109Val, p.Gly325Ser, and p.Ser461Cys) in our cohort had reduced total and V5-tagged EIF2AK2 protein expression. In comparison, the rare variant control *EIF2AK2* p.Leu114Gln, which we identified in a proband with discordant phenotype from our curation of the BG sequencing database, had no reduction in protein levels compared to *EIF2AK2* WT (**Fig3B, i-iii**). To examine whether the reduced EIF2AK2 protein stability associated with the p.Tyr133Phe, p.Ala109Val, p.Gly325Ser, and p.Ser461Cys variants was the result of impaired EIF2S1 signaling on general protein translation, we performed co-transfection with an unrelated protein, MAP1LC3A, tagged with GFP-RFP (GFP-RFP-MAP1LC3A). The co-transfection of *EIF2AK2* variants with GFP-RFP-MAP1LC3A had no effect on GFP-RFP-MAP1LC3A levels, indicating that overexpression of *EIF2AK2* variants in HEK293T or HeLa cells does not affect general protein translation (**Fig3B, ii**).

Based on our findings that HEK293T and HeLa cells are insensitive to EIF2AK2 overexpression or knockdown, we obtained three independent patient-derived fibroblast lines heterozygous for *EIF2AK2* p.Met11Leu, p.Tyr133Phe, and Ser461Cys. First, we examined the levels of EIF2S1 phosphorylation in the fibroblast lines and found a consistent reduction in p-EIF2S1 levels in all three lines (**Fig3C**). EIF2AK2 protein levels were stable in the heterozygous patient-derived skin fibroblast lines (**Fig3D**), indicating that the reduced p-EIF2S1 levels were likely due to impaired EIF2AK2 kinase activity. Next, we examined whether EIF2AK2 kinase activity can be stimulated in the heterozygous fibroblast lines by inducing cellular stress through the addition of polyinosinic:polycytidylic acid (poly-IC). Poly-IC is structurally similar to dsRNA, which is present in some viruses, and can activate the ISR pathway through EIF2S1 phosphorylation by EIF2AK family members^30^. Incubation with poly-IC activates Toll-like receptor 3 (TLR3), which recognizes dsRNA^31^, and the activated TLR3 recruits TRAF6, TAK1, and TAB2 to form the TAK1-complex^32–34^. EIF2AK2 is present in the poly-IC induced TAK1-complex and expression of a kinase inactive EIF2AK2 mutant protein inhibits poly-IC induction of the TLR3-mediated signaling pathway^32^. Furthermore, poly-IC stimulation of mammalian cells has been shown to upregulate the EIF2AK2-mediated phosphorylation of EIF2S1^10; 35; 36^. Together these findings suggest that poly-IC stimulation of mammalian cells triggers both a primary TLR3-mediated signaling event and a secondary EIF2Ak2-mediated signaling event following poly-IC uptake into cells^10^. Therefore, to test the functional consequences of the *EIF2AK2* variants we incubated the patient-derived skin fibroblasts with 10 μg/ml poly-IC for 24 hours and then assessed p-EIF2S1 levels by Western blot. Control (Ctrl) fibroblasts derived from unrelated healthy individuals show an increase in EIF2S1 phosphorylation upon addition of poly-IC (**Fig3E**). However, the fibroblast lines heterozygous for either *EIF2AK2* p.Met11Leu or p.Tyr133Phe failed to upregulate EIF2S1 phosphorylation in the presence of poly-IC (**Fig3E**). We were unable to test the poly-IC induction in the heterozygous *EIF2AK2* p.S461C fibroblasts as the line failed to expand after a few passages. Together, these results demonstrate that *EIF2AK2* p.Met11Leu, p.Tyr133Phe, and Ser461Cys impair the EIF2AK2 kinase activity required for EIF2S1 phosphorylation in fibroblasts.

The results of our clinical and molecular characterizations in mammalian cell lines and patient-derived fibroblasts show that *EIF2AK1* or *EIF2AK2* missense variants at key functional domains lead to neurodevelopmental disorders with overlapping symptoms. The *EIF2AK1* p.Ile448Val, *EIF2AK2* p.Met11Leu, *EIF2AK2* p.Tyr133Phe, and *EIF2AK2* p.Ser461Cys variants that we tested in either mammalian cell lines or patient-derived skin fibroblasts showed reduced kinase activity with impaired EIF2S1 phosphorylation.

Comparing genotypes and phenotypes within the cohort reveals several findings of interest. First, proband 1 with a *de novo EIF2AK1* p.Ile448Val variant has a distinct motor-predominant phenotype compared to the rest of the cohort with *de novo EIF2AK2* variants. Proband 1’s phenotype is primarily distinguished by motor developmental delay, speech articulation disorder, progressive spastic hemiplegia with hyper-reflexia, and age-appropriate cognition. The unrelated probands 2-8 have *de novo EIF2AK2* missense variants and their phenotypes are relatively more severe compared to proband 1. Common phenotypes in probands 2-8 include motor findings as well as ataxia, movement disorders, cognitive impairment, abnormal myelination, cerebral volume loss, reduced cerebellar vermis volume, and white matter findings. The LoF o/e score for *EIF2AK1* (0.47) is higher than for *EIF2AK2* (0.3), suggesting that *EIF2AK1* is more tolerant than *EIF2AK2* to LoF mutations. Therefore, the phenotypic spectrum associated with *EIF2AK1* variants may be milder than for *EIF2AK2* variants or there may be incomplete penetrance of *EIF2AK1* pathogenic variants. However, this determination is limited by the small sample size. Second, of the seven probands with *de novo EIF2AK2* missense variants, proband 2 is the only individual without a history of neurologic decompensation in the setting of fevers and illnesses. Although the reason for a relatively milder phenotype associated with the *EIF2AK2* p.Met11Leu variant remains unclear, it is possible the p.Met11Leu variant is less damaging as it did not reduce EIF2AK2 protein levels when overexpressed in mammalian cell lines.

Our functional data reveal that the *de novo* missense variants impair EIF2AK1 or EIFAK2 kinase activity and lead to reduced EIF2S1 phosphorylation. This impact on EIF2S1 activity would interfere with downstream molecular pathways critical for responding to cellular stressors. An abnormal stress response may underlie the neurologic decompensation and corresponding white matter alterations associated with fevers and illnesses in our cohort. Potential pathogenic mechanisms for these variants include gain-of-function, haploinsufficiency, and dominant-negative. A gain-of-function mechanism is less likely given the EIF2S1 phosphorylation and protein stability data, as well as the impaired response to poly-IC stimulation in fibroblasts. Haploinsufficiency may not be the sole contributor to the observed phenotypes in our probands given that LoF variants are present in gnomAD^24^, a family with thoracic aortic aneurysm syndrome was found to have a heterozygous deletion of chromosome 2p22.3-p22.2 involving *EIF2AK2* and 10 other genes^37^, and mouse models with constitutive loss of either *Eif2ak1* or *Eif2ak2* are viable and fertile without gross morphological abnormalities or neurologic findings^38; 39^. These findings are all consistent with *EIF2AK1* and *EIF2AK2* pLI scores of 0 and 0.06, respectively^24^, indicating that they are haplosufficient. Given that EIF2AK1 and EIF2AK2 require dimerization to phosphorylate their downstream target, a possible mechanism is that the *de novo* missense variants are dominant-negative mutations affecting the function of the wildtype protein.

The phosphorylation of EIF2S1 converts EIF2S1 into a competitive inhibitor of EIF2B, which activates the ISR^40^. Therefore, the impaired EIF2S1 phosphorylation we observed with the *de novo EIF2AK1* and *EIF2AK2* missense variants would likely impact the EIF2B-mediated regulation of the ISR. Pathogenic variants in any of the five genes encoding the subunits of the EIF2B protein complex (*EIF2B1, EIF2B2, EIF2B3, EIF2B4,* and *EIF2B5*) are associated with autosomal recessive childhood ataxia with central nervous system hypomyelination/vanishing white matter disease (CACH/VWM, MIM #603896)^41–45^. CACH/VWM is a chronic and progressive leukoencephalopathy characterized by neurologic decompensation in the setting of febrile illness and other stressors. Additional features of CACH/VWM include ataxia, spasticity, optic atrophy, epilepsy, loss of acquired developmental milestones, cognitive impairments, and coma^41; 46; 47^.

In conclusion, we have shown that pathogenic *EIF2AK1* and *EIF2AK2* missense variants cause a broad phenotypic spectrum including developmental delays, variable cognitive impairments, hypotonia, hypertonia, involuntary movements, ataxia, and white matter alterations. Individuals with *EIF2AK2* variants also exhibit sensitivity to febrile illness and commonly experience neurological regression, similar to CACH/VWM. The phenotypic overlap between CACH/VWM and our probands with *de novo* missense *EIF2AK1* and *EIF2AK2* variants suggest that deleterious missense variants in *EIF2AK1* and *EIF2AK2* cause an autosomal dominant neurodevelopmental syndrome that may share common pathogenic mechanisms with CACH/VWM disease.

## SUPPLEMENTAL DATA

Supplemental Data include Supplemental Materials and Methods.

## CONSORTIA

### Members of the Undiagnosed Diseases Network (UDN)

Maria T. Acosta, Margaret Adam, David R. Adams, Pankaj B. Agrawal, Mercedes E. Alejandro, Patrick Allard, Justin Alvey, Laura Amendola, Ashley Andrews, Euan A. Ashley, Mahshid S. Azamian, Carlos A. Bacino, Guney Bademci, Eva Baker, Ashok Balasubramanyam, Dustin Baldridge, Jim Bale, Michael Bamshad, Deborah Barbouth, Gabriel F. Batzli, Pinar Bayrak-Toydemir, Anita Beck, Alan H. Beggs, Gill Bejerano, Hugo J. Bellen, Jimmy Bennet, Beverly Berg-Rood, Raphael Bernier, Jonathan A. Bernstein, Gerard T. Berry, Anna Bican, Stephanie Bivona, Elizabeth Blue, John Bohnsack, Carsten Bonnenmann, Devon Bonner, Lorenzo Botto, Lauren C. Briere, Elly Brokamp, Elizabeth Burke, Lindsay C. Burrage, Manish J. Butte, Peter Byers, John Carey, Olveen Carrasquillo, Ta Chen Peter Chang, Sirisak Chanprasert, Hsiao-Tuan Chao, Gary D. Clark, Terra R. Coakley, Laurel A. Cobban, Joy D. Cogan, F. Sessions Cole, Heather A. Colley, Cynthia M. Cooper, Heidi Cope, William J. Craigen, Michael Cunningham, Precilla D'Souza, Hongzheng Dai, Surendra Dasari, Mariska Davids, Jyoti G. Dayal, Esteban C. Dell'Angelica, Shweta U. Dhar, Katrina Dipple, Daniel Doherty, Naghmeh Dorrani, Emilie D. Douine, David D. Draper, Laura Duncan, Dawn Earl, David J. Eckstein, Lisa T. Emrick, Christine M. Eng, Cecilia Esteves, Tyra Estwick, Liliana Fernandez, Carlos Ferreira, Elizabeth L. Fieg, Paul G. Fisher, Brent L. Fogel, Irman Forghani, Laure Fresard, William A. Gahl, Ian Glass, Rena A. Godfrey, Katie Golden-Grant, Alica M. Goldman, David B. Goldstein, Alana Grajewski, Catherine A. Groden, Andrea L. Gropman, Sihoun Hahn, Rizwan Hamid, Neil A. Hanchard, Nichole Hayes, Frances High, Anne Hing, Fuki M. Hisama, Ingrid A. Holm, Jason Hom, Martha Horike-Pyne, Alden Huang, Yong Huang, Rosario Isasi, Fariha Jamal, Gail P. Jarvik, Jeffrey Jarvik, Suman Jayadev, Yong-hui Jiang, Jean M. Johnston, Lefkothea Karaviti, Emily G. Kelley, Dana Kiley, Isaac S. Kohane, Jennefer N. Kohler, Deborah Krakow, Donna M. Krasnewich, Susan Korrick, Mary Koziura, Joel Krier, Seema R. Lalani, Byron Lam, Christina Lam, Brendan C. Lanpher, Ian R. Lanza, C. Christopher Lau, Kimberly LeBlanc, Brendan H. Lee, Hane Lee, Roy Levitt, Richard A. Lewis, Sharyn A. Lincoln, Pengfei Liu, Xue Zhong Liu, Nicola Longo, Sandra K. Loo, Joseph Loscalzo, Richard L. Maas, Ellen F. Macnamara, Calum A. MacRae, Valerie V. Maduro, Marta M. Majcherska, May Christine V. Malicdan, Laura A. Mamounas, Teri A. Manolio, Rong Mao, Kenneth Maravilla, Thomas C. Markello, Ronit Marom, Gabor Marth, Beth A. Martin, Martin G. Martin, Julian A. Martínez-Agosto, Shruti Marwaha, Jacob McCauley, Allyn McConkie-Rosell, Colleen E. McCormack, Alexa T. McCray, Heather Mefford, J. Lawrence Merritt, Matthew Might, Ghayda Mirzaa, Eva Morava-Kozicz, Paolo M. Moretti, Marie Morimoto, John J. Mulvihill, David R. Murdock, Avi Nath, Stan F. Nelson, John H. Newman, Sarah K. Nicholas, Deborah Nickerson, Donna Novacic, Devin Oglesbee, James P. Orengo, Laura Pace, Stephen Pak, J. Carl Pallais, Christina GS. Palmer, Jeanette C. Papp, Neil H. Parker, John A. Phillips III, Jennifer E. Posey, John H. Postlethwait, Lorraine Potocki, Barbara N. Pusey, Aaron Quinlan, Wendy Raskind, Archana N. Raja, Genecee Renteria, Chloe M. Reuter, Lynette Rives, Amy K. Robertson, Lance H. Rodan, Jill A. Rosenfeld, Robb K. Rowley, Maura Ruzhnikov, Ralph Sacco, Jacinda B. Sampson, Susan L. Samson, Mario Saporta, C. Ron Scott, Judy Schaechter, Timothy Schedl, Kelly Schoch, Daryl A. Scott, Lisa Shakachite, Prashant Sharma, Vandana Shashi, Jimann Shin, Rebecca Signer, Catherine H. Sillari, Edwin K. Silverman, Janet S. Sinsheimer, Kathy Sisco, Kevin S. Smith, Lilianna Solnica-Krezel, Rebecca C. Spillmann, Joan M. Stoler, Nicholas Stong, Jennifer A. Sullivan, Angela Sun, Shirley Sutton, David A. Sweetser, Virginia Sybert, Holly K. Tabor, Cecelia P. Tamburro, Queenie K.-G. Tan, Mustafa Tekin, Fred Telischi, Willa Thorson, Cynthia J. Tifft, Camilo Toro, Alyssa A. Tran, Tiina K. Urv, Matt Velinder, Dave Viskochil, Tiphanie P. Vogel, Colleen E. Wahl, Stephanie Wallace, Nicole M. Walley, Chris A. Walsh, Melissa Walker, Jennifer Wambach, Jijun Wan, Lee-kai Wang, Michael F. Wangler, Patricia A. Ward, Daniel Wegner, Mark Wener, Monte Westerfield, Matthew T. Wheeler, Anastasia L. Wise, Lynne A. Wolfe, Jeremy D. Woods, Shinya Yamamoto, John Yang, Amanda J. Yoon, Guoyun Yu, Diane B. Zastrow, Chunli Zhao, Stephan Zuchner

## ACKNOWLEDGEMENTS

The authors report no conflicts of interest. We thank the families and clinical staff at locations for participation in this study. We thank Seth Masters at the University of Melbourne, Australia for discussions and review of the manuscript. We thank Mingshan Xue, Fairouz Elsaeidi, Darrion Nguyen, and Maimuna Sali-Paul at BCM for critical review and feedback on the manuscript. Research reported in this manuscript was supported by the NIH Common Fund, through the Office of Strategic Coordination/Office of the NIH Director under Award Number(s) U01HG007709 (BCM clinical site), U01HG007708 (Stanford clinical site) and U01HG007942 (BCM sequencing core). The content is solely the responsibility of the authors and does not necessarily represent the official views of the National Institutes of Health. This was work was supported in part by U54NS093793 to H.J.B, by the Intramural Research Program of the National Human Genome Research Institute, and by the Common Fund, Office of the Director, NIH. H.J.B. is an investigator of the Howard Hughes Medical Institute. L.T.E, J.A.R, L.B, A.T., M.A., and H.T.C are supported in part by NIH grant U01HG007709. H.T.C.’s research effort was also supported by the American Academy of Neurology, Burroughs Wellcome Fund, and the Robert and Janice McNair Foundation. Sequencing and analysis for Proband 5 were provided by the Broad Institute of MIT and Harvard Center for Mendelian Genomics (Broad CMG) and was funded by the National Human Genome Research Institute, the National Eye Institute, and the National Heart, Lung and Blood Institute grant UM1 HG008900 and in part by National Human Genome Research Institute grant R01 HG009141.

## WEB RESOURCES

ENSEMBL VEP SIFT, http://useast.ensembl.org/info/docs/tools/vep/index.html; Exome Aggregation Consortium (EXAC), http://exac.broadinstitute.org/; Genome Aggregation Database (gnomAD), http://gnomad.broadinstitute.org/; Online Mendelian Inheritance in Man (OMIM), http://www.omim.org/; Model organism Aggregated Resources for Rare Variant ExpLoration (MARRVEL), http://marrvel.org/; Mutation Taster, http://www.mutationtaster.org; PolyPhen-2, http://genetics.bwh.harvard.edu/pph2/; Genic Intolerance (Residual Variation Intolerance Score), http://genic-intolerance.org

