## Supplemental Materials for "*De novo EIF2AK1* and *EIF2AK2* variants are associated with developmental delay, leukoencephalopathy, and neurologic decompensation"

### SUPPLEMENTAL EXPERIMENTAL PROCEDURES

#### Human subjects and sequencing studies

Informed consent for all subjects was obtained in accordance with research protocols that were approved by the institutional review board at Baylor College of Medicine, Stanford University, or at local institutions prior to testing. DNA was extracted for all subjects from peripheral blood mononuclear cells for trio exome sequencing (ES) in CLIA certified laboratories and variants were confirmed by Sanger sequencing.

For proband 1 and 7, trio ES was performed at GeneDX with Illumina SureSelect XT kit reagents and a HiSeq2500 platform (Illumina). For proband 2 and 6, trio ES was performed at Baylor Genetics through the Whole Genome Laboratory (https://www.bcm.edu/research/medical-genetics-labs/index.cfm?PMID=21319) using methods described[1]. Produced sequence reads were aligned to the GRCh37 (hg19) human genome reference assembly using the HGSC Mercury analysis pipeline (http://www.tinyurl.com/HGSC-Mercury/). Variants were determined and called using the Atlas2 suite to produce a variant call file[2].For the population comparisons we utilized data from the Exome Aggregation Consortium (ExAC), Cambridge, MA (http://exac.broadinstitute.org), Exome Variant Server, NHLBI GO Exome Sequencing Project (ESP), Seattle, WA (URL: http://evs.gs.washington.edu/EVS/), and Genome Aggregation Database (gnomAD), Cambridge, MA (http://gnomad.broadinstitute.org). Probands 1 and 2 were enrolled at the BCM UDN site. Baylor Genetics provided research reanalysis of proband 1's trio ES and UDN researchers used Codified Genomics for variant interpretation. For proband 3, trio ES was performed at Ambry Genetics with research analysis and variant interpretation by researchers at the Stanford UDN site. For proband 4, trio ES was performed at Children's Mercy Hospital Center for Pediatric Genomic Medicine as previously described[3]. For proband 5, trio ES was performed by the Center for Mendelian Genomics and the Broad Institute, analyzed the results with SEQR (https://seqr.broadinstitute.org) and VExP[4]. For proband 8, trio ES was conducted using genomic DNA from the proband and parents, the exonic regions and flanking splice junctions of the genome were captured using either the Clinical Research Exome v.2 kit (Agilent Technologies, Santa Clara, CA). Sequencing was done on a NextSeq500 Illumina system with 150bp paired-end reads. Reads were aligned to human genome build GRCh37/UCSC hg19, and analyzed for sequence variants using a custom-developed analysis tool[5]. Additional sequencing technology and variant interpretation protocol have been previously described[5]. Coverage on target for the index was ≥ 10x for 98.6% with a mean coverage of 200x.

All variant nomenclature uses GRCh37 (hg19) human genome reference assembly with GenBank: NM_014413.4 (*EIF2AK1*) and GenBank: NM_002759.3 (*EIF2AK2*).

**cDNA mutagenesis**

The cDNAs encoding the EIF2AK1/2 WT protein were subcloned from the respective Gateway donor vectors into the mammalian expression vector, pcDNA-DEST40 (with c-terminal V5 tag), via the Invitrogen Gateway LR Clonase II protocol. The stop codon was removed by mutagenesis to tag the protein with V5 tag. EIF2AK1/2 variants were introduced via site directed mutagenesis using the QuikChange II site-directed mutagenesis kit (Agilent, 200523) or NEB Q5 Site-Directed Mutagenesis Kit (E0554). Clones were sequenced and confirmed to be correctly subcloned into the pcDNA-DEST40 destination vector. Primers used for the mutagenesis are listed below:

| Primer | Sequence (5'-3') |
|---|---|
| EIF2AK1-A1342G-F | atcagggccatgaagaaaaacatttcttggcttcagatctc |
| EIF2AK1-A1342G-R | gagatctgaagccaagaaatgtttttcttcatggccctgat |
| EIF2AK2-A31T-F | ggtatgtattaagttcctccaagaagaaacctgctgaaagatc |

| | |
|---|---|
| EIF2AK2-A31T-R | gatctttcagcaggtttcttcttggaggaacttaatacatacc |
| EIF2AK2-A227G-F | gccttcttttccttactaagtatctcaacagctaatttggctg |
| EIF2AK2-A227G-R | cagccaaattagctgttgagatacttagtaaggaaaagaaggc |
| EIF2AK2-A227G-F | gccttcttttccttactaagtatctcaacagctaatttggctg |
| EIF2AK2-A227G-R | cagccaaattagctgttgagatacttagtaaggaaaagaaggc |
| EIF2AK2-C326T-F | ttacagttagtcttttcttctggacaattctattgataaggcctatg |
| EIF2AK2-C326T-R | cataggccttatcaatagaattgtccagaagaaaagactaactgtaa |
| EIF2AK2-T341A-F | cacactgttcataatttacagtttgtcttttcttctgggcaattcta |
| EIF2AK2-T341A-R | tagaattgcccagaagaaaagacaaactgtaaattatgaacagtgtg |
| EIF2AK2-A398T-F | ctgtcccattttgcatttaaaatgaaatccttctggccc |
| EIF2AK2-A398T-R | gggccagaaggatttcattttaaatgcaaaatgggacag |
| EIF2AK2-G973A-F | atcccaacagctattgtagtgaacaatatttacatgatcaagt |
| EIF2AK2-G973A-R | acttgatcatgtaaatattgttcactacaatagctgttgggatggatt |
| EIF2AK2-C1382G-F | atagtcttgcgaacaaatctgttctgggctcatgtatc |
| EIF2AK2-C1382G-R | gatacatgagcccagaacagatttgttcgcaagactat |

**Mammalian tissue culture**

HEK293T or HeLa cells were grown in high glucose Dulbecco's modified Eagle's medium (ThermoFisher Scientific, 11960) supplemented with 10% fetal bovine serum (Sigma, F0926) 1% (v:v) GlutaMAX (ThermoFisher Scientific, 35050061), and 1% (v:v) penicillin-streptomycin (GenDEPOT, CA005-010) and grown in a humidified incubator at 37°C with 5% $CO_2$.

**Poly-IC treatment**

Control or patient-derived skin fibroblasts were incubated in regular media with or without poly-IC (Sigma, P1530) (final concentration 10 μg/ml) for 24hrs. Cells were lysed for protein collection and Western blot analysis.

**DNA and siRNA Transfection**

Generation of mammalian expression vectors with EIF2AK1/2 WT or variants cDNAs are discussed above. GFP-RFP-MAP1LC3A tandem tagged constructs were provided by Marco

Sardiello (Baylor College of Medicine). siRNAs used for transient interference of EIF2AK1/2 are listed below:

| siRNA | Source | Catalog | Sequence (5'-3') |
|---|---|---|---|
| EIF2AK1-1 | Sigma | SASI_Hs01_00086018 | CGUUGUAUUUAGUAAGCCU |
| EIF2AK1-2 | Sigma | SASI_Hs01_00086021 | CCUUUACAAGACUUGUUAA |
| EIF2AK1-3 | ThermoFisher | s25822 | GACGGAAAGACUUACGUUAtt |
| EIF2AK2-1 | Sigma | SASI_Hs01_00019634 | GAGGUUUACAUUUCAAGUU |
| EIF2AK2-2 | Sigma | SASI_Hs01_00019640 | GUCAGAAGCAGGGAGUAGU |
| EIF2AK2-3 | ThermoFisher | s11185 | GAUUAAGGGUGCAACUAAAtt |

For DNA and siRNA transfection, HEK293T or HeLa cells were seeded in 6-well plates till 60% confluence. To introduce siRNA, cells are transfected with scramble or EIF2AK1/2 siRNAs using Lipofectamine RNAiMAX for 3 days using standard protocol. To introduce DNA, cells are transfected with mammalian expression vectors using Lipofectamine 3000 for 2 days following standard protocol. The transfection ratio of DNA (μg) to Lipofectamine 3000 (μl) was 1 to 2. Transfected cells were lysed with 300 μl of Lysis Buffer (2% SDS, 50mM Tris, 2mM EDTA).

**Protein sample collection and Western blotting**

To collect protein samples, cultured cells were washed with PBS once, and then lysed with Lysis Buffer (2% SDS, 50mM Tris, 2mM EDTA). Protein concentration was measured and normalized with Pierce™ BCA Protein Assay Kit (ThermoFisher, Catalog No. 23225). The whole cell lysates were fractioned by SDS-PAGE (Bio-Rad 4–20% precast polyacrylamide gel) and transferred to nitrocellulose membranes using tank electroblotting transfer system according to the manufacturer's protocols (Bio-Rad). Nitrocellulose membranes were then incubated with 5% BSA in TBST (10 mM Tris, pH7.4, 150mM NaCl, 0.1% Tween-20) before incubating with primary antibodies. Primary antibodies used for Western blotting are listed below:

| Antibody | Dilution | Source | Catalog |
|---|---|---|---|
| V5 | 1:5000 | Invitrogen | R960-25 |
| EIF2AK2 | 1:1000 | Cell Signaling | 12297S |
| GAPDH | 1:5000 | Cell Signaling | 2118S |
| Phospho-EIF2S1 | 1:1000 | Cell Signaling | 3597S |
| GFP | 1:2000 | Invitrogen | A11122 |
| EIF2S1 | 1:2000 | abcam | ab26197 |
| ACTIN | 1:5000 | MP Biomedicals | ICN691001 |

**Western blot image collection and analysis**

Western blot images were acquired using a Bio-Rad ChemiDoc™ Imaging Systems and all images were collected by the imaging system within the linear range. Densitometric measurement of the bands were performed with ImageJ. Statistical analysis of the quantification was performed with R. We first confirmed the normality of the data ($p = 0.05$, Shapiro-Wilk test), and then used Student's t-test to measure the difference between groups. Data shown as mean ± standard error of mean. Number of replicates, n, quantified per test group is stated in the figure.
